# Long-term transcriptional activity at zero growth by a cosmopolitan rare biosphere member

**DOI:** 10.1101/284430

**Authors:** Bela Hausmann, Claus Pelikan, Thomas Rattei, Alexander Loy, Michael Pester

## Abstract

Microbial diversity in the environment is mainly concealed within the rare biosphere (all species with <0.1% relative abundance). While dormancy explains a low-abundance state very well, the mechanisms leading to rare but active microorganisms remain elusive. We used environmental systems biology to genomically and transcriptionally characterise *Candidatus* Desulfosporosinus infrequens, a low-abundance sulfate reducer cosmopolitan to freshwater wetlands, where it contributes to cryptic sulfur cycling. We obtained its near-complete genome by metagenomics of acidic peat soil. In addition, we analyzed anoxic peat soil incubated under *in situ*-like conditions for 50 days by *Desulfosporosinus*-targeted qPCR and metatranscriptomics. The *Desulfosporosinus* population stayed at a constant low abundance under all incubation conditions, averaging 1.2 × 10^6^ 16S rRNA gene copies per cm^3^ soil. In contrast, transcriptional activity of *Ca*. D. infrequens increased at day 36 by 56- to 188-fold when minor amendments of acetate, propionate, lactate, or butyrate were provided with sulfate, as compared to the no-substrate-control. Overall transcriptional activity was driven by expression of genes encoding ribosomal proteins, energy metabolism and stress response but not by expression of genes encoding cell growth-associated processes. Since our results ruled out growth of these highly active microorganisms, they had to invest their sole energy for maintenance, most likely counterbalancing acidic pH conditions. This finding explains how a rare biosphere member can contribute to a biogeochemically relevant process while remaining in a zero growth state.

## Importance

The microbial rare biosphere represents the largest pool of biodiversity on Earth and constitutes, in sum of all its members, a considerable part of a habitat’s biomass. Dormancy or starvation are typically used to explain the persistence of low-abundance microorganisms in the environment. We show that a low-abundance microorganism can be highly transcriptionally active while remaining in a zero growth-state over prolonged time periods. Our results provide evidence that this zero-growth at high-activity state is driven by maintenance requirements. We show that this is true for a microbial keystone species, in particular a cosmopolitan but permanently low-abundance sulfate reducer in wetlands that is involved in counterbalancing greenhouse gas emission. In summary, our results provide an important step forward in understanding rare biosphere members relevant for ecosystem functions.

## Introduction

The vast majority of microbial diversity worldwide is represented by the rare biosphere (1–4). This entity of microorganisms consists of all microbial species that have an arbitrarily defined relative population size of <0.1% in a given habitat at a given time (1–4). The rare biosphere is opposed by a much smaller number of moderately abundant or very abundant microbial species (≥0.1% and ≥1.0% relative abundance, respectively) (5), which are thought to be responsible for the major carbon and energy flow through a habitat as based on their cumulative biomass. However, there is accumulating experimental evidence that the rare biosphere is not just a “seed bank” of microorganisms that are waiting to become active and numerically dominant upon environmental change (3, 6), but also harbors metabolically active microorganisms with important ecosystem functions (4).

First hints for metabolically active rare biosphere members were evident from seasonal patterns of marine bacterioplankton species. Here, many taxa that displayed recurring annual abundance changes were of low abundance and even during their bloom periods never reached numerically abundant population sizes (7–9). In soil environments, removal of low-abundance species by dilution-to-extinction had a positive effect on intruding species, suggesting that active low-abundance species pre-occupy ecological niches and thus slow down invasion (10–12). Soil microorganisms of low relative abundance were also shown to play a role in community-wide species interactions, e.g, by being involved in the production of antifungal compounds that protect plants from pathogens (13) or by constituting the core of microorganisms that respond to the presence of a particular plant species (14). Other examples include microorganisms with a specialized metabolism that sustain stable low-abundance populations in an ecosystem (3). For example, N_2_-fixing microorganisms in the ocean (15) or sulfate-reducing microorganisms (SRM) in peatlands (5, 16, 17) were shown to fulfill such gatekeeper functions.

A peatland *Desulfosporosinus* species was one of the first examples identified as an active rare biosphere member contributing to an important ecosystem function (16). This SRM is involved in the cryptic sulfur cycle of peatlands (5, 16), which in turn controls the emission of the greenhouse gas CH_4_ from these globally relevant environments (17). Although porewater sulfate concentrations are typically quite low in peatlands (<300 μM) (17), these environments are characterized by temporally fluctuating high sulfate reduction rates (up to 1800 nmol cm^−3^ day^−1^) (17). These rates can be in the same range as in sulfate-rich marine surface sediments, where sulfate reduction is one of the major anaerobic carbon degradation pathways (18, 19). In low-sulfate peatlands, such high sulfate reduction rates can only be maintained by rapid aerobic or anaerobic re-oxidation of reduced sulfur species back to sulfate (17). Since SRM generally outcompete methanogens and syntrophically associated fermenters (20), they exert an important intrinsic control function on peatland CH_4_ production (21–23). This is important, since natural wetlands, such as peatlands, are estimated to be responsible for 30% of the annual emission of this potent greenhouse gas (24–26).

Little is known about the ecophysiology of metabolically active but low-abundance microorganisms. This lack of knowledge is clearly founded in their low numerical abundance making it inherently difficult to study their metabolic responses or even to retrieve their genomes directly from the environment. In a preceding study, we could show that the low-abundance peatland *Desulfosporosinus* species mentioned above follows an ecological strategy to increase its cellular ribosome content while maintaining a stable low-abundance population size when exposed to favorable, sulfate-reducing conditions (5). This was unexpected since metabolic activity in bacteria and archaea is typically followed by growth (in terms of cell division or biomass increase) if they are not severely energy or nutrient limited (27) or engaged in major maintenance processes coping with (environmental) stress (28). The studied *Desulfosporosinus* species is found worldwide in a wide range of low-sulfate wetlands including peatlands, permafrost soils, and rice paddy fields (5). This emphasizes its importance as a model organism for active rare biosphere members. In this study, we used an environmental systems biology approach to deepen our understanding of the ecophysiology of this rare biosphere member. In particular, we retrieved its genome by metagenomics from native and incubated peat soil and followed its transcriptional responses in peat soil microcosms, which were exposed to different environmental triggers that mimicked diverse *in situ* conditions.

## Materials and Methods

### Genome assembly, binning, and phylogenetic inference

Sampling of peat soil from the acidic peatland Schlöppnerbrunnen II (Germany), DNA-stable isotope probing (DNA-SIP), total nucleic acids extraction, metagenome sequencing and assembly, and coverage-based binning was described previously (5, 16, 29). In brief, DNA from native peat soil (10-20 cm depth) and DNA pooled from 16 ^13^C-enriched fractions (density 1.715-1.726 g mL^−1^) of a previous DNA-SIP experiment with soil from the same site (16) was sequenced using the Illumina HiSeq 2000 system. DNA-SIP was performed after a 73-day incubation (again 10-20 cm depth) that was periodically amended with small dosages of sulfate and first a mixture of unlabeled formate, acetate, propionate, and lactate for two weeks and thereafter a mixture of ^13^C-labeled formate, acetate, propionate, and lactate (all in the lower µM-range) (16). Raw reads were quality filtered, trimmed, and co-assembled (native soil: 385 million reads; DNA-SIP: 576 million reads) using the CLC Genomics Workbench 5.5.1 (CLC Bio). Differential coverage binning was applied to extract the *Desulfosporosinus* metagenome-assembled genome (MAG) (30). As expected (16), the *Desulfosporosinus* MAG was of low abundance in the native soil with an average coverage of 0.026 while enriched in the SIP sample with an average coverage of 34 (detailed per scaffold in Table S2). A side effect of sequencing a DNA-SIP sample is an apparent G+C content skew, which was normalized arbitrarily to improve binning using the following formula (29, 31):

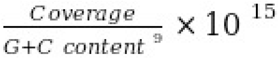

Scaffolds encoding the 16S and 23S rRNA genes were successfully identified using paired-end linkage data (30). Completeness, contamination, and strain heterogeneity was estimated using CheckM 1.0.6 (32).

Phylogenomic analysis of the *Desulfosporosinus* MAG was based on a concatenated set of 34 phylogenetically informative marker genes as defined by (32) and the Bayesian phylogeny inference method PhyloBayes using the CAT-GTR model (33). 16S rRNA gene-based phylogeny was inferred using the ARB SILVA database r126 as a reference (34), the SINA aligner (35), and the substitution model testing and maximum likelihood treeing method IQ-TREE (36). Pairwise 16S rRNA gene sequence identities were calculated with T-Coffee 11 (37). Pairwise average nucleic and amino acid identities (ANI, AAI) (38) between protein-coding genes of the *Desulfosporosinus* MAG and reference genomes were calculated as described previously (29)

### Genome annotation

The genome was annotated using the MicroScope annotation platform (39). Annotation refinement for selected genes was done as follows: proteins with an amino acid identity ≥40% (over ≥80% of the sequence) to a Swiss-Prot entry (40), curated MaGe annotation (39), or protein described in the literature were annotated as true homologos of known proteins. The same was true, if classification according to InterPro families (41, 42), TIGRFAMs (43), and/or FIGfams (44) led to an unambiguous annotation. Proteins with an amino acid identity ≥25% (over ≥80% of the sequence) to a Swiss-Prot or TrEMBL (40) entry were annotated as putative homologs of the respective database entries. In addition, classification according to COG (45) or InterPro superfamilies, domains, or binding sites were used to call putative homologs in cases of an unambiguous annotation. Membership to syntenic regions (operons) was considered as additional support to call true or putative homologs.

### Metatranscriptomics and quantitative PCR from single-substrate incubations

For this study, we re-analysed qPCR and metatranscriptomic data sets of of anoxic peat soil slurry microcosms that were described previously under different aspects (5, 29). In brief, anoxic microcosms were incubated at 14 °C in the dark for 50 days and regularly amended with either low amounts of sulfate (76-387 μM final concentrations) or incubated without an external electron acceptor. Formate, acetate, propionate, lactate, butyrate (<200 μM), or no external electron donor was added to biological triplicates each. DNA and/or RNA were extracted from the native soil and after 5, 8, 15, 26, 36, and 50 days of incubations. Quantitative PCR data describing 16S rRNA gene copies of the complete *Desulfosporosinus* population in comparison to the overall bacterial and archaeal community (5) was re-analyzed to put the metatranscriptome data into the perspective of population dynamics. PCR conditions are given in (5). Metatranscriptome sequencing was done from each of the biological replicates using the Illumina HiSeq 2000/2500 platform (27-188 million reads per sample). Raw reads were quality-filtered as described previously (29) and mapped to the *Desulfosporosinus* MAG in a background of all other metagenome-assembled scaffolds using Bowtie 2 at default settings (46). Counting of mapped reads to protein-coding genes (CDS) was performed with featureCounts 1.5.0 (47).

### Statistical analysis of *Desulfosporosinus-specific* transcripts

Counts of mapped transcript reads were normalized to the length of the respective gene and the sequencing depth of the respective metatranscriptome, resulting in FPKM (fragments per kilobase per million total fragments) values. Thereafter, we used an unsupervised approach to identify CDS expression stimulated by sulfate and the different substrates regimes. First, we applied the DESeq2 R package (48, 49) to identify differentially expressed CDS. Treatments without external sulfate added and samples after 8 days of incubations had too little transcript counts to be used for a statistical approach. Therefore, we limited our analysis to pairwise comparison of sulfate-stimulated microcosms after 36 days of incubations. We compared each substrate regime to the no-substrate controls and each other. The set of all significantly differentially expressed CDS (FDR-adjusted *p*-value < 0.05) were further clustered into response groups. For clustering, we calculated pairwise Pearson’s correlation coefficients (*r*) of variance stabilized counts (cor function in R), transformed this into distances (1−*r*), followed by hierarchical clustering (hclust function in R). Variance stabilisation was performed using the rlog function of the DESeq2 package. Spearman’s rank correlation of FPKM values for each gene to the total relative mRNA counts was performed with cor.test in R using the data from all treatments and replicates.

### Sequence data availability

The MAG SbF1 is available at MicroScope (https://www.genoscope.cns.fr/agc/microscope/) and is also deposited under the GenBank accession number OMOF01000000. Metagenome and - transcriptomic data is available at the Joint Genome Institute (https://genome.jgi.doe.gov/) and are also deposited under the GenBank accession numbers PRJNA412436 and PRJNA412438, respectively.

## Results

### A near complete genome of a rare biosphere member from peat soil

We obtained the population genome of the low-abundance *Desulfosporosinus* species by coassembly and differential coverage binning of metagenomes obtained from native peat soil and ^13^C-labelled fractions of a DNA-stable isotope probing experiment of the same peatland (Fig. S1) (29). The high quality metagenome-assembled genome (MAG) SbF1 had a size of 5.3 Mbp (on 971 scaffolds), a G+C content of 42.6%, a checkM-estimated completeness of 98.0%, a potential residual contamination of 3.9%, and 10% strain heterogeneity. Besides 16S and 23S rRNA genes, SbF1 carried 6440 protein-coding genes (CDS), five 5S rRNA gene copies, 59 tRNAs, and 37 other ncRNAs, making a total of 6543 predicted genomic features. The genome size and G+C content was in the same range as observed for genomes of cultured *Desulfosporosinus* species (3.0-5.9 Mbp and 42-44%, respectively) (50–54). Scaffolds encoding rRNA genes had a higher coverage compared to the average coverage of all scaffolds (Fig. S1), indicating multiple *rrn* operon copies, as is known from genomes of other *Desulfosporosinus* species (on average 9.3 *rrn* operons, range: 8-11) (55).

16S rRNA-based phylogenetic tree reconstruction placed SbF1 into a well supported clade together with *Desulfosporosinus* sp. 44a-T3a (98.3% sequence identity), *Desulfosporosinus* sp. OT (98.8%), and *Desulfosporosinus* sp. 5apy (98.1%). The most similar validly described species was *Desulfosporosinus lacus* with a sequence identity of 97.5% (Fig. S2a). Phylogenomics confirmed *Desulfosporosinus* sp. OT as the closest relative (Fig. S2b) with average amino and nucleic acid identities (AAI and ANI) of 77% and 79%, respectively (Fig. S3). The intra-genus AAI variability of *Desulfosporosinus* species was 69-93% (Fig. S3). Therefore, MAG SbF1 represents a novel species in this genus based on species-level thresholds of 99% for the 16S rRNA gene (56) and 96.5% for ANI (38).

### The versatile energy metabolism of the low-abundance *Desulfosporosinus*

*Desulfosporosinus* sp. MAG SbF1 encoded the complete canonical pathway for dissimilatory sulfate reduction (Fig. 1, Table S1). This encompassed the sulfate adenylyltransferase (Sat), adenylyl-sulfate reductase (AprBA), dissimilatory sulfite reductase (DsrAB), and the sulfide- releasing DsrC, which are sequentially involved in the reduction of sulfate to sulfide. In addition, genes encoding the electron-transferring QmoAB and DsrMKJOP complexes were detected, with their subunit composition being typical for *Desulfosporosinus* species (50, 51, 53, 54). Other *dsr* genes included *dsrD, dsrN*, and *dsrT* (57) with hitherto unvalidated function, *fdxD*, which encodes a [4Fe4S]-ferredoxin, and a second set of DsrMK-family encoding genes (*dsrM2* and *dsrK2).* SbF1 also encoded the trimeric dissimilatory sulfite reductase AsrABC (anaerobic sulfite reductase) (58).

**Fig. 1.**
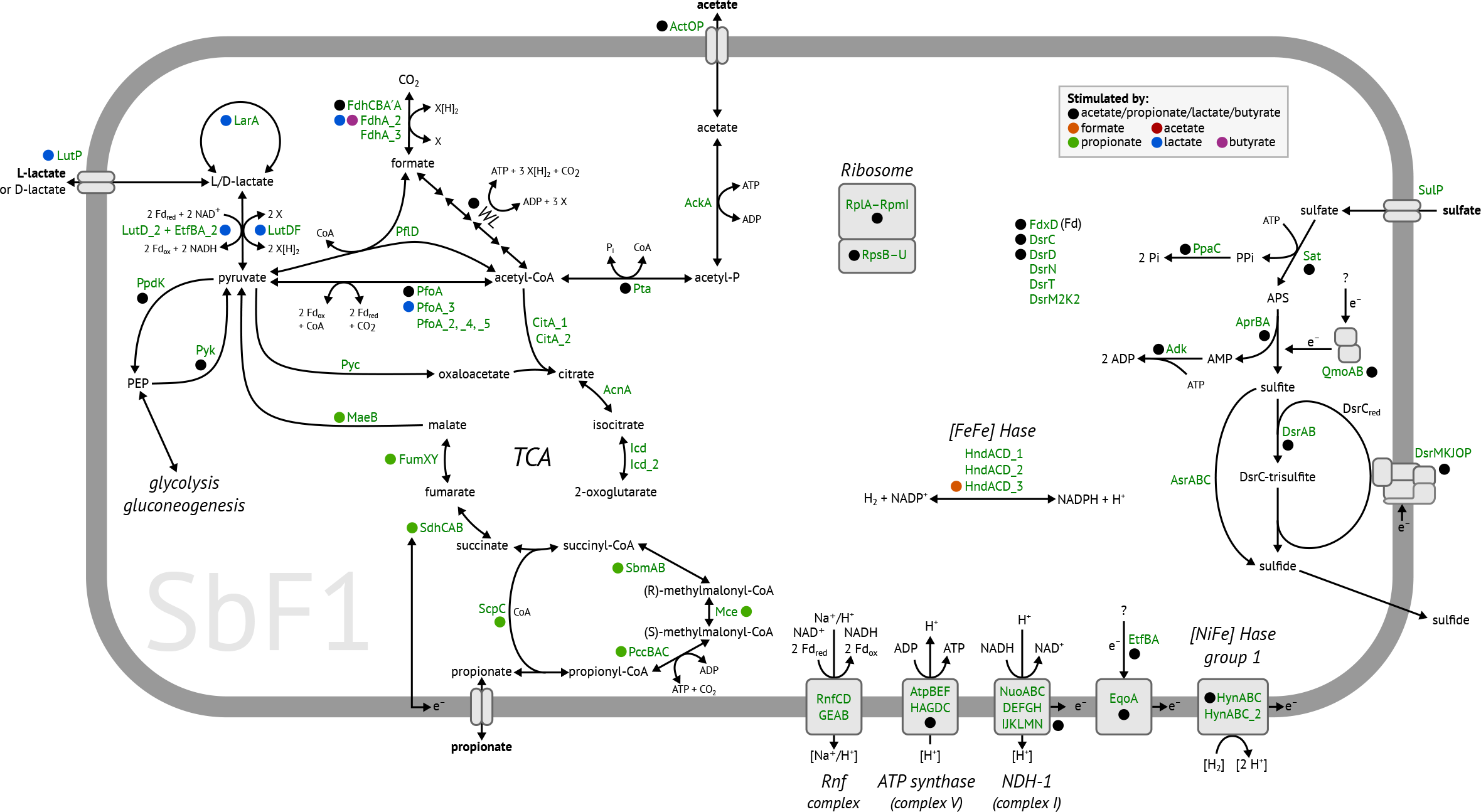
Metabolic model of *Desulfosporosinus* sp. MAG SbF1. Gene expression stimulated by specific substrates in combination with sulfate is indicated by coloured points. Paralogous genes are indicated by an underscore followed by a number. Plus signs indicates proposed protein complexes. Details for all genes are given in Table S1 and transcription patterns are shown in Fig. 4. For the citric acid cycle and anaplerotic reactions, carriers of reducing equivalents and further by-products are not shown. The following abbreviations were used. X: unknown reducing equivalent carrier, e.g., NAD^+^ or ferredoxin. WL: Wood-Ljungdahl pathway consisting of enzymes encoded by the *acs* operon, MetF, FolD, FchA, and Fhs. TCA: citric acid cycle. FDH: formate dehydrogenase. Hase: hydrogenase. NDH-1: NADH dehydrogenase 1. LDH: lactate dehydrogenase.

SbF1 carried genes for both complete and incomplete oxidation of propionate and lactate. In addition, the ability to utilize acetate, formate, or H2 as electron donors was encoded (Fig. 1). All enzymes necessary for propionate oxidation to the central metabolite pyruvate (including those belonging to a partial citric acid cycle) were encoded on two scaffolds (Table S1). For lactate utilization, SbF1 carried three paralogs of glycolate/D-lactate/L-lactate dehydrogenase family genes. However, the substrate specificity of the encoded enzymes could not be inferred from sequence information alone. The transcription of *lutDF* and *lutD_2* was stimulated by the addition of L-lactate (Fig. 1), which indicates that these genes encode functional lactate dehydrogenases (LDH). The third paralog (*glcDF*, Table S1) was not stimulated by lactate. LutDF was organised in an operon with a lactate permease (LutP) and a lactate regulatory gene (*lutR).* LutD_2 was organised in a operon with an electron-transferring flavoprotein (EtfBA_2), which resembled the electron-confurcating LDH/Etf complex in *Acetobacterium woodii* (59). LDHs have been shown to utilize both L- and D-lactate (59, 60). However, SbF1 also encoded a lactate racemase (LarA) and a lactate racemase-activating system (LarEBC) for interconversion of both stereoisomers (61).

Pyruvate, the intermediate product in propionate and lactate degradation, can be further oxidized to acetyl-CoA with either one of several pyruvate-ferredoxin oxidoreductases (PfoA) or formate C-acetyltransferase (PflD). Acetyl-CoA can then be completely oxidized to CO_2_ via the Wood-Ljungdahl pathway (62), which is complete in SbF1 (Fig. 1, Table S1) and present in the genomes of all other sequenced *Desulfosporosinus* species (50, 51, 53, 54). Alternatively, acetyl-CoA may be incompletely oxidized to acetate via acetyl-phosphate by phosphate acetyltransferase (Pta) and acetate kinase (AckA). Pta and AckA are bidirectional enzymes, opening the possibility that acetate could be degraded via these two enzymes and the downstream Wood-Ljungdahl pathway to CO_2_.

Formate and H_2_ represented additional potential electron donors for SbF1. Its genome encoded three formate dehydrogenases (FDH). FDH-1 consists of three subunits (*fdhCBA*) while FDH-2 (FdhA_2) and FDH-3 (FdhA_3) are monomeric enzymes. In addition, [NiFe] hydrogenases of group 1 and 4f, as well as [FeFe] hydrogenases of group A (63) were encoded. Homologs of genes for butyrate oxidation were missing in SbF1 (64), which is in contrast to other *Desulfosporosinus* species (e.g., *Desulfosporosinus* orientis). Both glycolysis and gluconeogenesis were complete. However, neither a glucokinase or a phosphotransferase system was found (PTS). Coupling of electron transfer to energy conservation could be mediated in SbF1 by a H^+^/Na^+^-pumping Rnf complex (RnfCDGEAB) (65) and a NADH dehydrogenase (respiratory complex I, NuoABCDEFGHIJKLMN). In addition, the complete gene set for ATP synthase (AtpABCDEFGH) was identified (Fig. 1, Table S1).

### Long-term transcriptional activity of *Desulfosporosinus* sp. MAG SbF1 at zero growth

Naturally occuring hot spots of sulfate reducing activity in peat soil (66–69) were mimicked by periodically amending sulfate in the lower µM-range to anoxic peat microcosms (every 3-7 days) and comparing this to unamended (i.e., methanogenic) control microcosms. In addition, sulfate reducing and methanogenic microcosms received, in triplicates, periodic amendments of either formate, acetate, propionate, lactate, or butyrate as compared to controls without amendment. Substrate supply did generally not exceed 100-200 μM thus again mimicking *in situ* concentrations of these naturally occurring organic carbon degradation intermediates in peatlands (5). The overall *Desulfosporosinus* population remained stable throughout the incubation period in the various microcosms (on average 1.2 × 10^6^ 16S rRNA gene copies per cm^3^ of soil, Fig. 2a). Compared to the total bacterial and archaeal community, this resembled a relative abundance of 0.018% when corrected for the average 9.3 *rrn* operons per genome in the genus *Desulfosporosinus* (55). The 16S rRNA gene of *Desulfosporosinus* sp. MAG SbF1 was 100% identical to OTU0051, which dominated the *Desulfosporosinus* population as evident from a previously published 16S rRNA (gene) amplicon survey of the same microcosms (5). In contrast to its stable low-abundance, the overall *Desulfosporosinus* population substantially increased its 16S rRNA copy numbers by 2.2, 4.9, 5.9, or 13.6-fold in sulfate reducing incubations stimulated by either acetate, propionate, lactate, or butyrate, respectively. In contrast, *Desulfosporosinus* 16S rRNA copy numbers remained stable or even slightly decreased in the sulfate-amended no-substrate-control and the methanogenic incubations (Fig. S4). Again, these increases were mainly reflected in changes of OTU0051 (*Desulfosporosinus* sp. MAG SbF1) as shown in the amplicon study mentioned above (5).

**Fig. 2.**
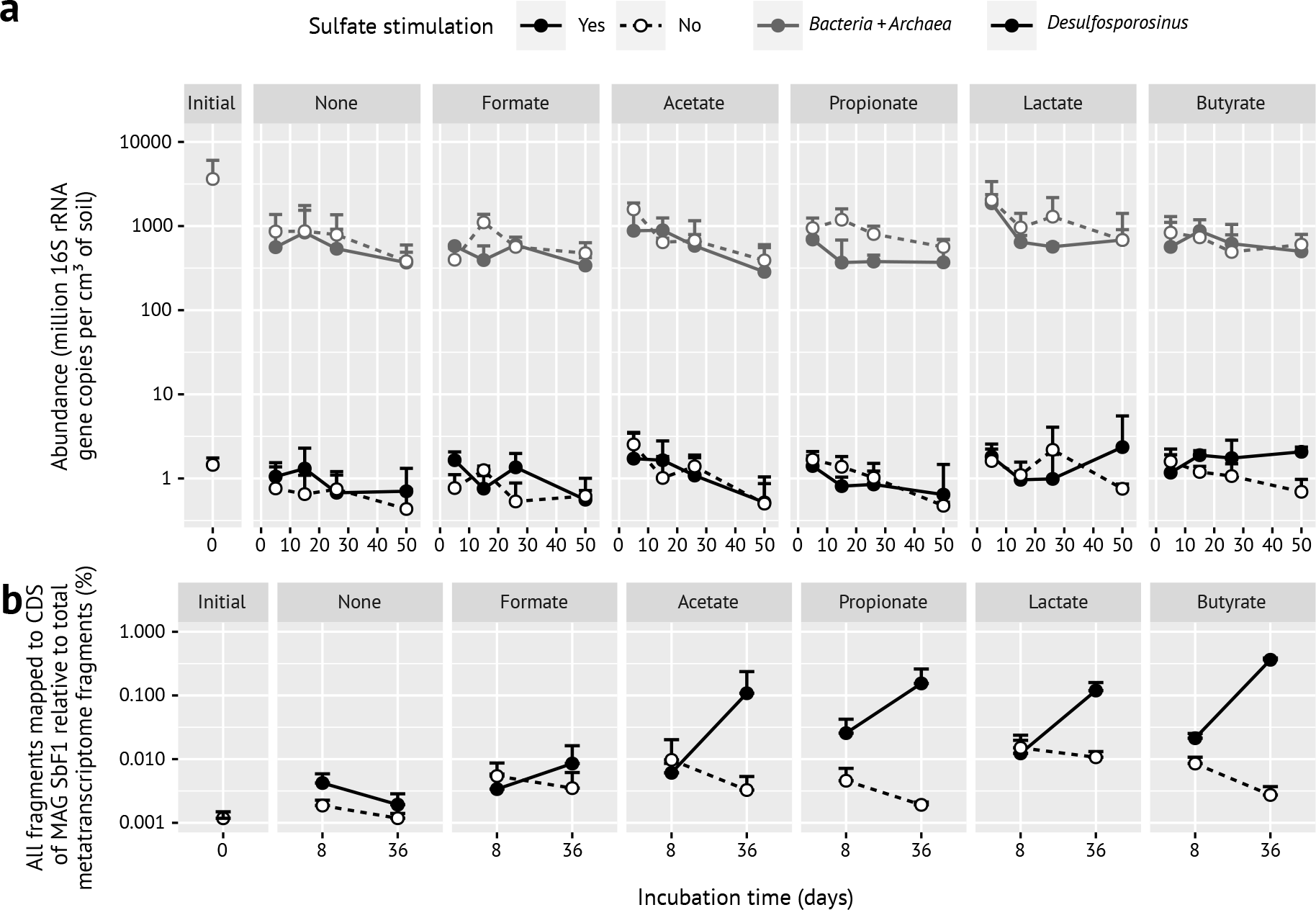
(a) Time-resolved absolute abundance of the *Desulfosporosinus* population (in black) as compared to all Bacteria and *Archaea* (in grey) in anoxic peat soil microcosms under various in- situ like conditions as determined by quantitative PCR (modified from (5)). Error bars represent one standard deviation of the mean (n=3; n=2 for propionate with sulfate stimulation, all days, and butyrate with sulfate stimulation, day 50). (b) Corresponding overall transcriptional changes (mRNA of all CDS) of *Desulfosporosinus* sp. MAG SbF1 in the same anoxic microcosms. Error bars represent one standard deviation of the mean (n=3; n=2 for propionate with sulfate stimulation).

We used metatranscriptomics of the same microcosms to analyse whether this strong increase in 16S rRNA copies at zero growth was accompanied by gene expression of metabolic pathways and cell-growth associated processes in *Desulfosporosinus* sp. MAG SbF1. Compared to the initial soil, the overall transcriptional activity of SbF1 steadily increased at day 8 and 36 in sulfate reducing incubations stimulated by either acetate, propionate, lactate, or butyrate. In contrast, all methanogenic incubations as well as the sulfate reducing formate and nosubstrate incubations showed after an initial stimulation until day 8, a steady or even mildly decreasing overall transcriptional activity (Fig. 2b). At day 36, normalized mRNA counts of SbF1 were 56-, 80-, 62-, or 188-fold higher in sulfate reducing incubations stimulated by either acetate, propionate, lactate, or butyrate, respectively, as compared to the no-substrate-control and constituted between 0.11 ± 0.13% (acetate) and 0.36 ± 0.02% (butyrate) of all transcripts in the corresponding metatranscriptomes (Fig. 2b). This substrate-specific activity was driven by the increased transcription of genes encoding ribosomal proteins as general activity markers (Fig. 3, Table S1) and energy metabolism genes including all canonical dissimilatory sulfate reduction genes (Fig. 4, Table S1). For example, Spearman’s rank correlation coefficients of normalised *dsrA* and *dsrB* transcript counts as compared to the sum of normalised SFb1 mRNA counts were 0.91 and 0.90, respectively (FDR-adjusted *p*-value < 0.001). Normalised transcript counts of other enzyme complexes involved in the central metabolism of SbF1 such as the ATP synthase, the NADH dehydrogenase (complex I), and ribosomal proteins followed the same transcriptional pattern (Fig. 4, Table S1) with an average Spearman’s rank correlation coefficients of 0.79 ± 0.07 (n = 72, FDR-adjusted *p*-value < 0.05) to the sum of normalised SFb1 mRNA counts. Interestingly, transcription of genes encoding proteins involved in general stress response were stimulated as well. In particular, genes encoding the universal stress promotor UspA, the GroES/GroEL and DnaK chaperons, and the proteolytic subunit of ATP-dependent Clp protease (ClpP) showed an increased transcription (Fig. 4) with an average Spearman’s rank correlation coefficients of 0.76 ± 0.04 (n = 5, FDR- adjusted *p*-value < 0.05) to the sum of normalised SFb1 mRNA counts.

**Fig. 3.**
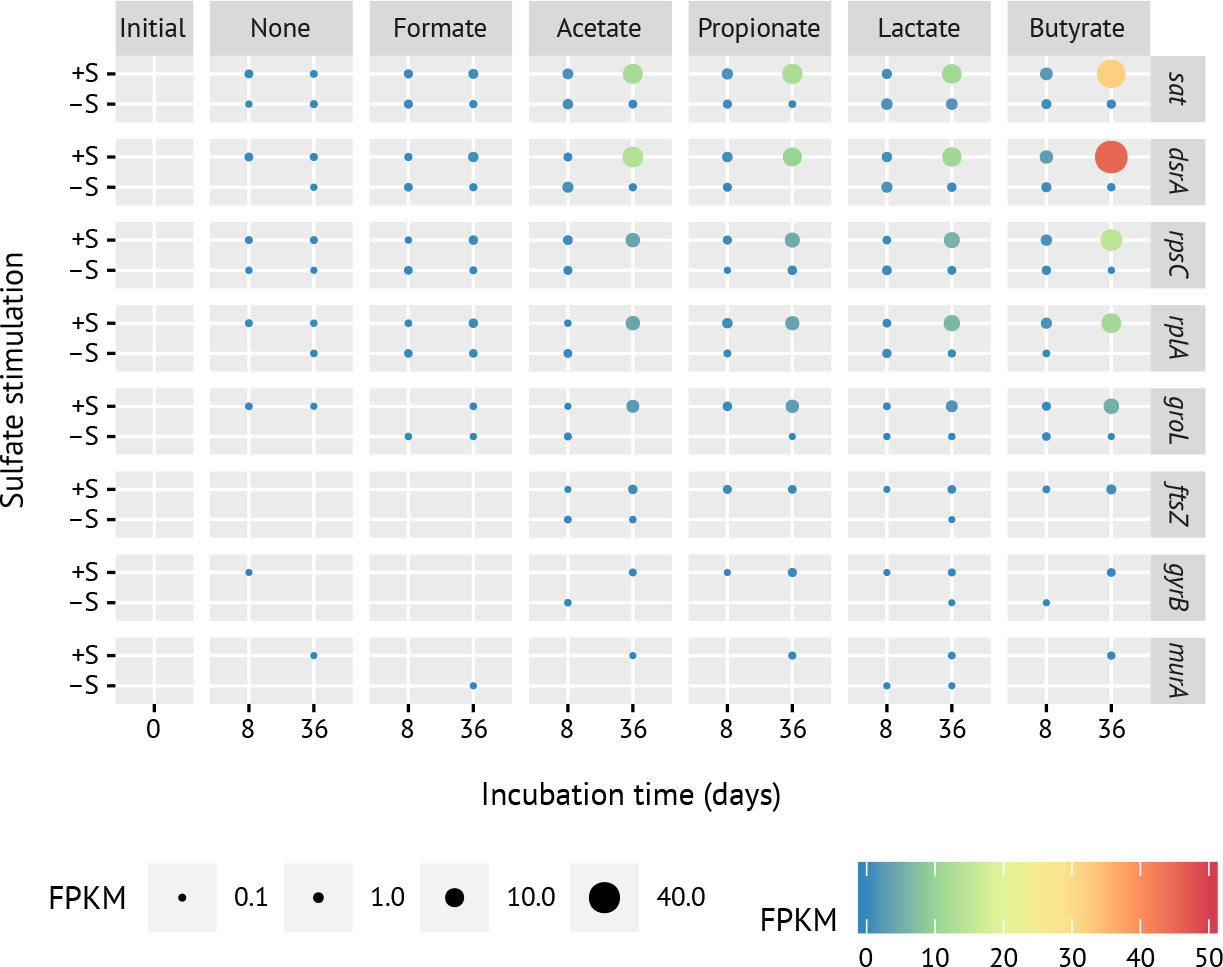
Time-resolved transcriptional changes of selected genes representing the sulfate-reduction pathway (*sat, dsrA*), ribosomal proteins of the large (*rplA*) and small subunit (*rpsC*), cell division (*ftsZ*), DNA replication (*gyrB*), and peptidoglycan synthesis (*murA).* Panels represent the various substrate incubations: initial, initial peat soil to set up peat microcosms; +/−S, incubations with or without external sulfate. The size and color of the dots represent average FPKM values of the respective normalised gene expression.

**Fig. 4.**
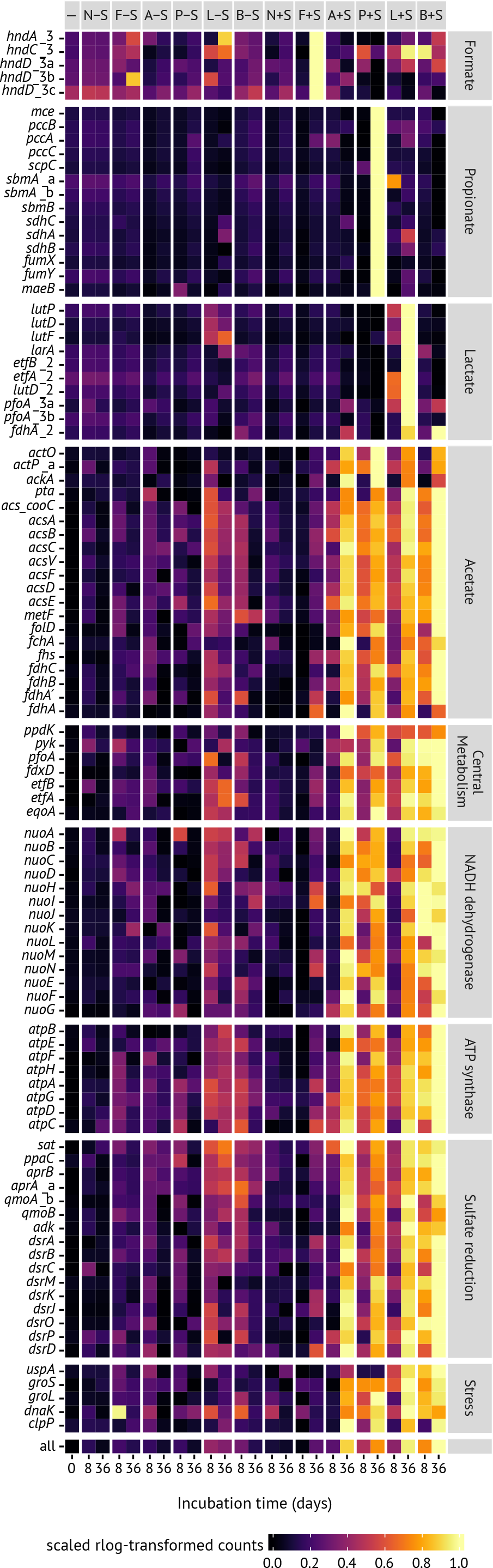
Transcription patterns of whole pathways and central enzyme complexes involved in the carbon and energy metabolism of *Desulfosporosinus* sp. MAG SbF1 under *in* situ-like conditions. In addition, transcription patterns of general stress response proteins are shown. Mean abundance for the native soil (—) and each incubation treatment and time point is shown. Supplemented substrates are indicated by initials and addition of external sulfate is depicted by −S/+S (columns). Abundance values are normalised variance-stabilised counts x, which were scaled from 0 to 1 for each CDS using the formula [x - min(x)] / max[x - min(x)]. Incompletely assembled genes are indicated by _a, _b, and _c.

To evaluate whether a hidden turnover of biomass (cryptic growth) was underlying the stable *Desulfosporosinus* population, we screened COG categories D, L, and M for expression of indicator genes that encode functions in cell division (e.g., *ftsZ* or *minE*), DNA replication (e.g., *gyrBA, dnaC*, and *dnaG*), and cell envelope biogenesis (e.g., *murABCDEFGI*), respectively. Genes that unambiguously encoded such functions (Table S1) showed only very minor or no increases in transcripts over time (Fig. 3, detailed in Fig. S5). Extension of this analysis to all genes belonging to COG D (n = 73), L (n = 280), and M (n = 215) showed that the average Spearman’s rank correlation coefficients to the sum of normalised mRNA counts was only 0.45 ± 0.13 (FDR-adjusted *p*-value < 0.05, Table S1).

We also analysed genes reported to be upregulated immediately after phage infection, which is an important ecological control of bacterial population size. Respective genes in *Bacillus subtilis* encode, e.g., functions in DNA and protein metabolism and include the ribonucleoside- diphosphate reductase (*nrdEF*) and aspartyl/glutamyl-tRNA amidotransferase (*gatCAB*) (70). However, homologs in SbF1 did not show increased expression in the incubations with increased total transcriptional activity (Table S1). This was reflected in an average Spearman’s rank correlation coefficient of only 0.60 ± 0.06 (n = 4, FDR-adjusted *p*-value < 0.05) to the sum of normalised SFb1 mRNA counts. The same was true when screening for active sporulation of a *Desulfosporosinus* subpopulation as an alternative explanation for a stable low-abundance population. The identified sporulation genes (*spo0A-spoVT*) did not show any prominent increase in transcript numbers as well, with the only exception of *spoIIIAD*, which was stimulated in propionate- and sulfate-amended microcosms (Table S1). Again, expression of genes involved in sporulation had a low average Spearman’s rank correlation coefficient of 0.44 ± 0.13 (n = 22, FDR-adjusted *p*-value < 0.05) to the sum of normalised SFb1 mRNA counts.

The individual incubation regimes triggered in addition transcriptional activation of the respective substrate degradation pathways of *Desulfosporosinus* sp. MAG SbF1. For example, all genes necessary for the conversion of propionate to pyruvate were overexpressed only upon addition of propionate and sulfate but not in any other incubation type. The same was true for lactate degradation, where genes encoding the lactate permease, lactate racemase and two of the detected lactate dehydrogenases were overexpressed upon addition of both lactate and sulfate, but not in incubations with lactate only (Fig. 4). Although genes encoding phosphotransacetylase and acetate kinase were overexpressed under lactate and propionate, the complete Wood-Ljungdahl pathway was overexpressed as well, which indicates that at least part of these substrates were completely degraded to CO_2_ rather than to acetate and CO_2_. This conclusion was supported by the overexpression of the Wood-Ljungdahl pathway in incubations amended with acetate and sulfate. Interestingly, the Wood-Ljungdahl pathway was also overexpressed upon addition of butyrate and sulfate. Under such conditions, *Desulfosporosinus* sp. MAG SbF1 apparently relies on a synthrophic lifestyle based on acetate uptake as it lacked the capability for butyrate oxidation; albeit failed recovery of the butyrate degradation pathway during binning cannot be excluded.

## Discussion

Current knowledge on the interconnection of energy metabolism, gene expression, cell division, and population growth in microorganisms is mainly based on pure cultures that are maintained in the laboratory. Under ideal conditions, a single *Escherichia coli* cell would grow to a population with the mass of the Earth within 2 days. Clearly, this does not occur but the discrepancy between potential and actual growth underscores that microorganisms spend the vast majority of their time not dividing (27, 71). A large fraction of these microorganisms is part of the rare biosphere. For example, in the studied peatland, the sum of all low-abundance species made up approximately 12% of the total bacterial and archaeal 16S rRNA genes (5). In other soils, low-abundance *Alphaproteobacteria* and *Bacteroidetes* alone constituted in sum 10% and 9% of the total bacterial population, respectively, while all low-abundance populations summed up to 37% of all bacteria (14). Upon strong environmental change, low-abundance microorganisms often grow to numerically abundant populations and replace dominant species as observed for microbial community changes after an oil spill (72, 73) or in the response of soil microorganisms towards the presence of plants (14). However, subtle environmental changes (5) or recurring seasonal shifts (7, 9, 74) often lead to rather small shifts in low- abundance populations without rare biosphere members becoming numerically dominant.

The low-abundance *Desulfosporosinus* sp. MAG SbF1 represents an interesting case of the latter response type. When exposed to favorable, sulfate-reducing conditions in peat soil microcosms, the overall *Desulfosporosinus* population did not increase its population size of about 1.2 × 10^6^ 16S rRNA gene copies cm^−3^ soil (Fig. 2a) but strongly increased its cellular ribosome content by up to one order of magnitude (Fig. S4) (5). In a preceding 16S rRNA (gene) amplicon study which analysed the same microcosms, we could show that *Desulfosporosinus* OTU0051 is the major constituent of this *Desulfosporosinus* population and correlated best in its 16S rRNA response to sulfate turnover among all identified SRM (5). Here, we re-analyzed these microcosms to expand upon this observation by genome-centric metatranscriptomics and to test whether the increase in cellular ribosome content is indeed translated into transcriptional and, as a consequence, metabolic activity. *Desulfosporosinus* OTU0051 was 100% identical to the 16S rRNA gene of *Desulfosporosinus* sp. MAG SbF1, which was retrieved in this study and as such represented the major *Desulfosporosinus* population. In support of this conclusion, increases in 16S rRNA copies of the overall *Desulfosporosinus* population (Fig. S4) clearly corresponded to increased transcription of genes coding for ribosomal proteins in *Desulfosporosinus* sp. MAG SbF1 (Fig. 3, Table S1) (5). This cellular ribosome increase under sulfate-reducing conditions correlated well to an increase in all normalised mRNA counts (Fig. 2b). This is the first time that changes in population-wide 16S rRNA levels are proven to be directly linked to transcriptional activity for a rare biosphere member.

Analyzing the transcriptional response of a rare biosphere member under *in* situ-like conditions opens the unique opportunity to gain insights into its ecophysiology. *Desulfosporosinus* sp. MAG SbF1 clearly overexpressed its sulfate reduction pathway under sulfate amendment when supplied with either acetate, lactate, propionate, or butyrate as compared to the no-substrate and the methanogenic controls (Fig. 4). Detailed analysis of the transcribed carbon degradation pathways showed that *Desulfosporosinus* sp. MAG SbF1 is able to oxidise propionate, lactate, and acetate completely to CO_2_. Under butyrate-amended conditions, it presumably relied on syntrophic oxidation of acetate supplied by a primary butyrate oxidiser. This shows that *Desulfosporosinus* sp. MAG SbF1 is capable of utilising diverse substrates that represent the most important carbon degradation intermediates measured in peatlands (75, 76). Such a generalist lifestyle is of clear advantage in peat soil given the highly variable nutrient and redox conditions (75, 76). These fluctuations are caused by the periodically changing water table that steadily shifts the oxic-anoxic interface (67, 77). In addition, the complex flow paths of water create distinct spatial and temporal patterns (hot spots and hot moments) of various biogeochemical parameters including sulfate and substrate availability, to which peat microorganisms have to adapt (66–69).

The question remains, which mechanisms are at work that keep the transcriptionally active *Desulfosporosinus* sp. MAG SbF1 population in a stable low-abundance state? Ongoing growth could be hidden by continuous predation, viral lysis, or active sporulation of a major subpopulation. To answer this question, we analysed expression patterns of genes involved in cell growth-associated processes. Compared to the strong overexpression of metabolic or ribosomal protein genes, transcription of genes essential for DNA replication, cell division, and cell envelope biogenesis did not increase or only marginally (Fig. 3, Fig. S5). In contrast, retentostat studies on cultured *Firmicutes* held in a (near)-zero growth state revealed that expression of genes involved in cell growth, central energy metabolism, and the translational apparatus were always co-regulated, either showing a joint increased expression in *Bacillus subtilis* (78) or an invariable expression in Lactobacillus plantarum (79) when comparing active growth to (near)-zero growth. In addition, there is experimental evidence that in the lag phase of batch cultures, i.e., in the transition from no growth to growth, transcription of growth- related genes is not stable but increases due to the overall activation of cellular processes (80). In this context, the lack of an increasing transcription of growth-related genes would clearly indicate a state of (near-)zero growth rather than an actively dividing population that is kept stable by an equally high growth and mortality or sporulation rate. This conclusion is further supported by the lack of overexpressed sporulation genes or genes upregulated directly after phage attack (Table S1; Table S3).

Nevertheless, the ATP generated by the induced energy metabolism has to be consumed. If not used for growth, it has to be invested completely for maintenance according to the Herbert-Pirt relation q_s_ = m_s_ + μ/Y_sx_^max^, where q_s_ is the biomass-specific consumption rate, m_s_ is the maintenance coefficient, µ is the specific growth rate, and Y_sx_^max^ is the the maximum growth yield (81, 82). Based on the the concept of a species-independent maintenance energy requirement as laid out by (83), and further developed by (28), it can be calculated that *Desulfosporosinus* sp. MAG SbF1 would need to consume 2.1 fmol sulfate per day to maintain a single cell in our incubations when, e.g., incompletely oxidizing lactate to acetate (detailed in Supplementary Information). This is in agreement with experimentally determined maintenance requirements of *Desulfotomaculum putei* (84), but two orders of magnitude smaller than the cell-specific sulfate reduction rates of *Desulfosporosinus* sp. MAG SbF1 estimated previously in a similar experimental setup of the same peat soil by (16) (here the responsive but low-abundance *Desulfosporosinus* OTU was 99.8% identical to the 16S rRNA gene of *Desulfosporosinus* sp. MAG SbF1). However, maintenance requirements are known to increase upon production of storage compounds or to counterbalance environmental stress (28). We found no indication for the former scenario but observed overexpression of the universal stress promotor UspA, which is one of the most abundant proteins in growth-arrested cells (85). In addition, we observed overexpression of the chaperons GroES/GroEL and DnaK and of the protease ClpP, which were all previously linked to low pH stress response at the expense of ATP consumption (86–90). Since the pH in the analyzed peat soil incubations varied between 4.1-5.0 (5), coping with a low pH would be the most parsimonious explanation for increased maintenance requirements. In this context, one may speculate whether the overexpressed ATP synthase might have operated as an ATPase to pump protons out of the cell at the expense of ATP hydrolysis, which is a known response mechanisms towards mildly acidic pH (90). Similar, the overexpressed sulfate reduction pathway including complex I and the membrane quinone shuttle might have been co-utilized as proton pump without harvesting the membrane potential for ATP generation. Since active sulfate reduction would also consume protons in the vicinity of *Desulfosporosinus* sp. MAG SbF1 and thus slowly increase its surrounding pH, a high metabolic activity at concomitant zero growth controlled by maintenance requirements would make sense.

Our results are important in the context of the increasing awareness that the microbial rare biosphere is not only the largest pool of biodiversity on Earth (1–4) but in sum of all its low- abundance members constitutes also a large part of the biomass in a given habitat (5, 14). Understanding the mechanisms governing this low-abundance prevalence and its direct impact on ecosystem functions and biogeochemical cycling is thus of utmost importance. *Desulfosporosinus* sp. MAG SbF1 has been repeatedly shown to be involved in cryptic sulfur cycling in peatlands (5, 16) — a process that counterbalances the emission of the greenhouse gas methane due to the competitive advantage of SRM as compared to microorganisms involved in the methanogenic degradation pathways (20). This species can be found worldwide in low-sulfate environments impacted by cryptic sulfur cycling including not only peatlands but also permafrost soils, rice paddies, and other wetland types (5). Here, we provided proof that *Desulfosporosinus* sp. MAG SbF1 is indeed involved in the degradation of important anaerobic carbon degradation intermediates in peatlands while sustaining a low-abundance population. It has a generalist lifestyle in respect to the usable carbon sources, re-emphasizing its importance in the carbon and sulfur cycle of peatlands. Our results provide an important step forward in understanding the microbial ecology of biogeochemically relevant microorganisms and show that low-abundance keystone species can be studied "in the wild" using modern environmental systems biology approaches.

### Proposal of *Candidatus* Desulfosporosinus infrequens

Based on its phylogenetic placement and novel ecophysiological behaviour, we propose that *Desulfosporosinus* sp. MAG SbF1 represents a novel species with the provisional name *Candidatus* Desulfosporosinus infrequens sp. nov. (in.fre’quens. L. adj. *infrequens*, rare, referring to its low relative abundance). Based on its genome-derived metabolic potential and support from metatranscriptomics, *Ca.* D. infrequens is capable of complete oxidation of acetate, propionate and lactate with sulfate as the electron acceptor, with further potential for oxidation of molecular hydrogen (Fig. 1).

## Acknowledgements

We are grateful to Mads Albertsen, Norbert Bittner, Tijana Glavina del Rio, Florian Goldenberg, Craig Herbold, Stephan Köstlbacher, Per H. Nielsen, Ulrich Stingl, and Susannah G. Tringe for sequence analysis and technical support. We further thank Bernhard Schink for help in naming *Ca.* D. infrequens, Kenneth Wasmund for valuable feedback, and Johannes Wittmann, Jan-Ulrich Kreft, and Silvia Bulgheresi for helpful expert opinions. We acknowledge the LABGeM (CEA/IG/Genoscope & CNRS UMR8030) and the France Génomique National infrastructure (funded as part of Investissement d’avenir program managed by Agence Nationale pour la Recherche, contract ANR-10-INBS-09) for support with the MicroScope annotation platform. The work conducted by the Joint Genome Institute was supported by the Office of Science of the U.S. Department of Energy under Contract No. DE-AC02-05CH11231. This research was financially supported by the Austrian Science Fund (FWF, P23117-B17 to MP and AL, P25111- B22 to AL), the U.S. Department of Energy (CSP605 to MP and AL), the German Research Foundation (DFG, PE 2147/1-1 to MP), and the European Union (FP7-People-2013-CIG, Grant No PCIG14-GA-2013-630188 to MP).

## Supplementary Information

### Supplementary Methods

#### Calculation of minimum sulfate turnover for maintenance

The minimum sulfate turnover required for maintenance was calculated according to the species-independent Arrhenius equation outlined in (1). Here, m_e_ = Ae^−Ea/RT^ with m_e_ as the ree energy consumption rate for zero growth, A as a constant factor for anaerobic microorganisms (4.99 × 10^12^ kJ g d.wt.^−1^ d^−1^), E_a_ as constant activation energy (69.4 kJ mol^−1^), R as the universal gas constant (8.314 J mol^−1^ K^−1^), and T as temperature in K. We used a temperature of 14 °C (288.15 K) for our calculations because this was the temperature at which the incubations were performed. The resulting m_e_ was converted to cell-specific sulfate reduction rates required for maintenance based on the energy yield of a sulfate reducer when converting lactate to acetate (−160.4 kJ mol sulfate^−1^) (2) and a conversion factor of dry weight biomass to cell numbers of 2.9 × 10^13^ g d.wt. cell^−1^ (3).

#### Supplementary Tables

##### Table S1

Summary of all genomic features in *Desulfosporosinus* sp. MAG SbF1. Genes encoding the energy metabolism or central cellular functions are given first. COG class IDs were assigned by MaGe (Cognitor, www.ncbi.nlm.nih.gov/COG/). bactNOG and NOG IDs were assigned by best- match principle (4, 5). Spearman’s rank correlation is given for each gene’s normalized transcript counts as compared to the sum of normalized mRNA counts (FDR-adjusted *p*-values are indicated by asterisks: *, < 0.05; **, < 0.01; ***, < 0.001). Expression clusters represent the clusters assigned by correlation and hierarchical clustering analysis. The next five columns are log2 fold-changes of expression levels after 36 days of incubation in the sulfate-stimulated microcosms (i.e., substrate vs no-substrate-control). Missing fold-changes are due to all counts being zero in both compared treatments. Ranks are based on mean fragments per kilobase per million total fragments (FPKM). Also here, only data of sulfate- stimulated microcosms after 36 days of incubation are shown in addition to the native soil. Missing ranks indicate that expression was never detected in any replicate. Fragmented, i.e., mainly incompletely assembled genes are indicated by _a, _b, and _c. A ^1^ or ^2^ in the strand column indicates that this CDS is either the first or last on a scaffold, respectively (depending on the reading frame).

##### Table S2

Characteristics and coverage of all scaffolds belonging to *Desulfosporosinus* sp. MAG SbF1. The two scaffolds with the highest coverage encode the 23S and 16S rRNA genes, respectively.

##### Table S3

Expression levels of selected CDS in the analysed anoxic peat soil microcosms given in FPKM (mean ± one standard deviation). Loci are sorted as in Table S1. Headers display the individual treatments used in the peat soil microcosms: without and with external sulfate added; amended substrate; and days of incubation.

#### Supplementary Figures

**Fig. S1.**
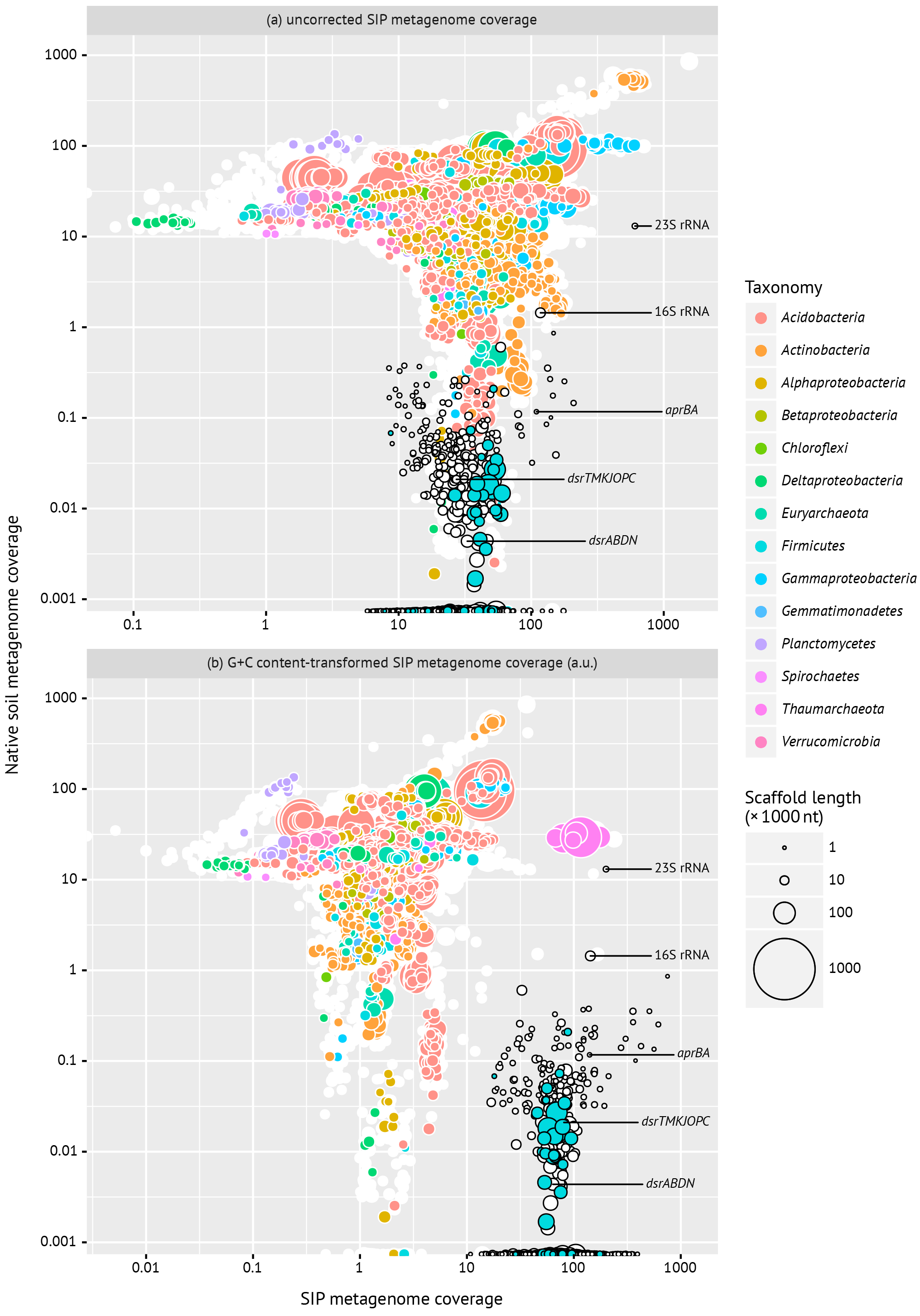
Differential coverage plots of assembled scaffolds with *Desulfosporosinus* sp. MAG SbF1 scaffolds highlighted by black circles. The average coverage per scaffold in the SIP metagenome is visualized without (a) and with (b) G+C content transformation (see Materials and Methods). Taxonomic affiliation is indicated by color and based on BLAST as described previously (6). White circles represent unclassified scaffolds. Only scaffolds >10 000 nt length are shown, except when belonging to SbF1. Scaffolds encoding selected genes in SbF1 are labelled accordingly.

**Fig. S2.**
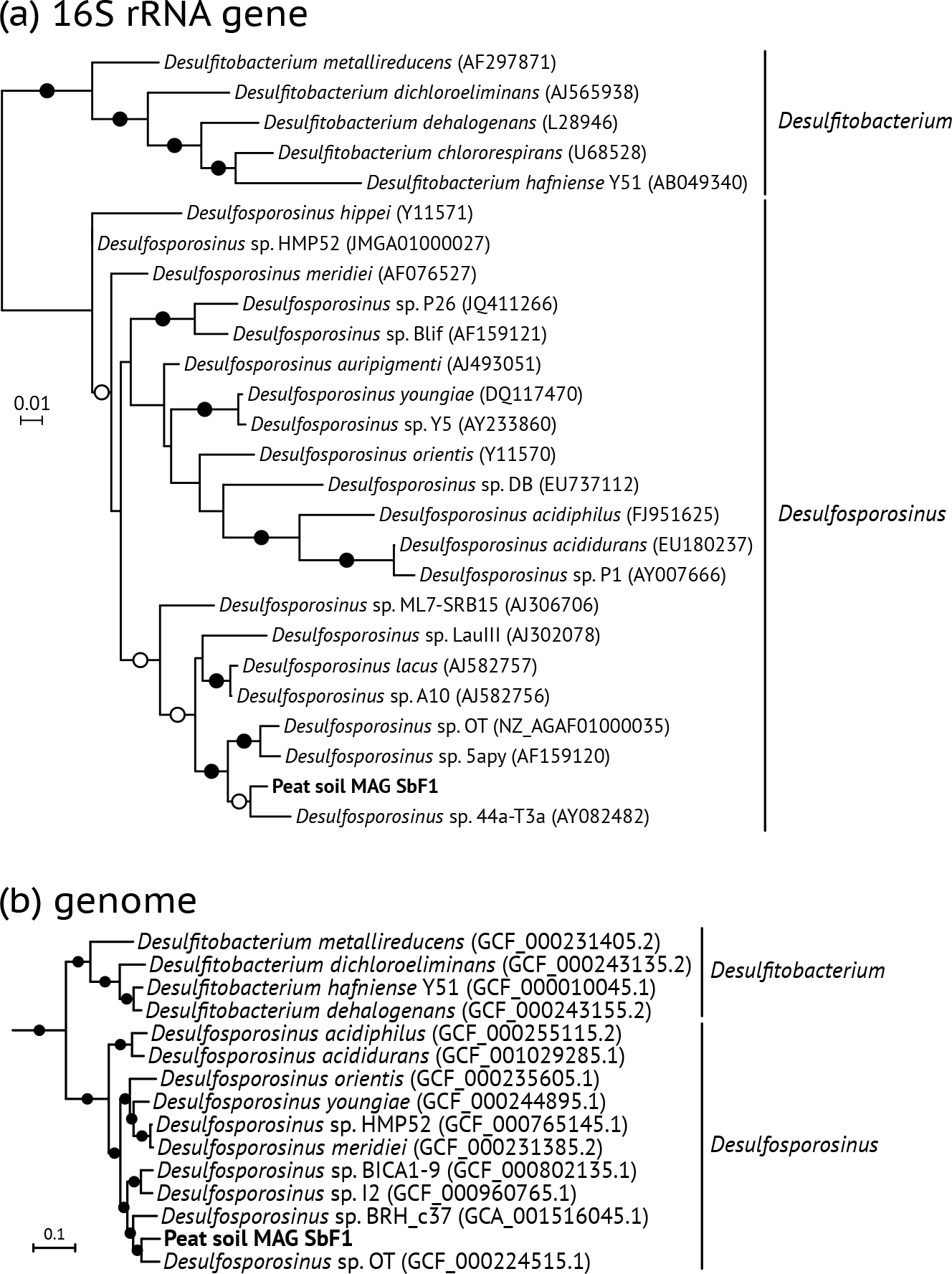
(a) Maximum likelihood 16S rRNA gene tree of species belonging to the genera *Desulfosporosinus* and *Desulfitobacterium*. Branch supports of ≥0.9 and ≥0.7 are indicated by filled and open circles, respectively. GenBank accession numbers are given in parentheses. (b) Bayesian inference phylogenomic tree showing the phylogenetic placement of *Desulfosporosinus* sp. MAG SbF1. All branches were supported >0.9 (filled circles). The tree was rooted against genomes from the *Acidobacteria, Proteobacteria*, and *Verrucomicrobia* (not shown). Genome accession numbers are given in parentheses.

**Fig. S3.**
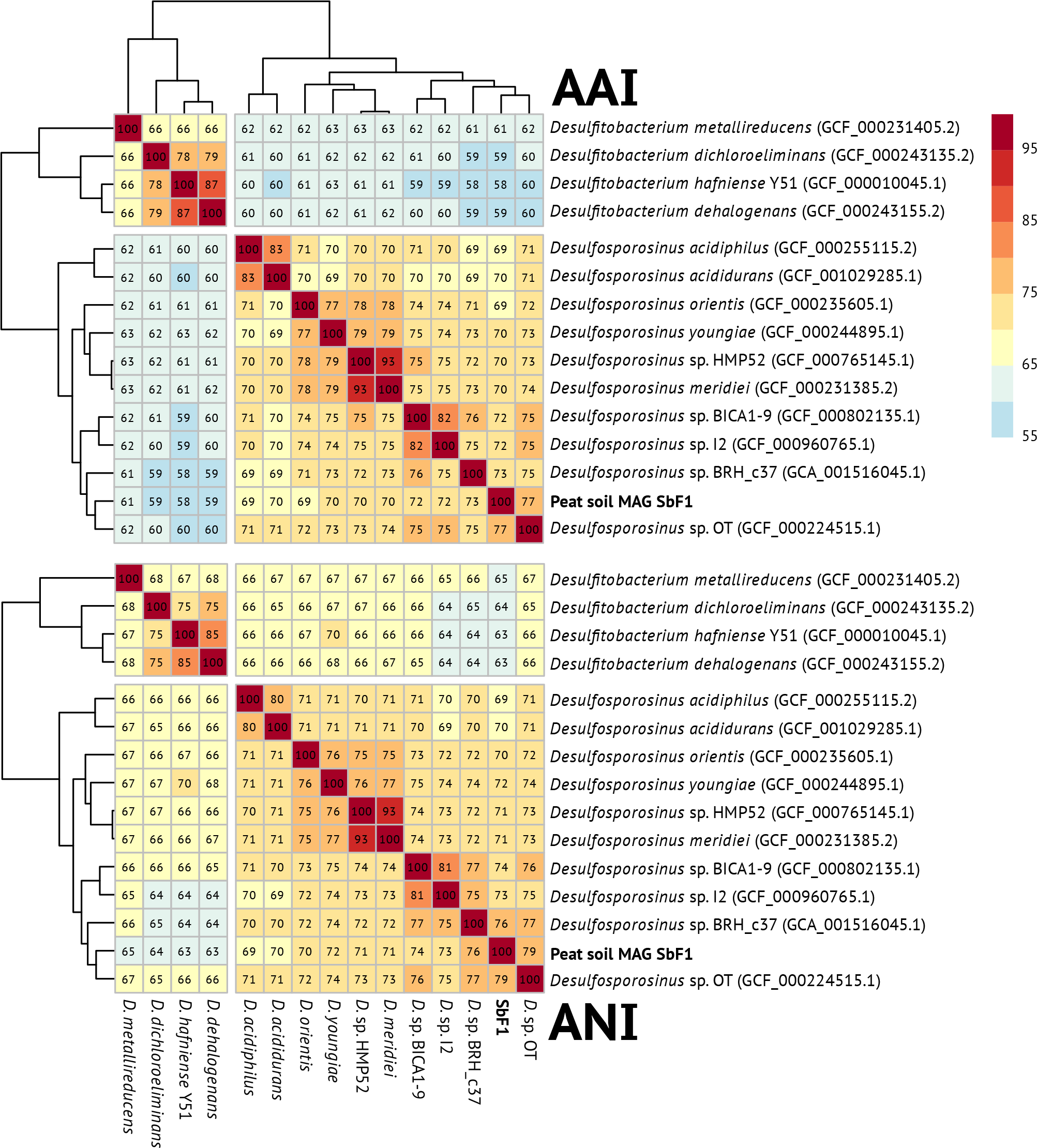
Two-way average amino and nucleic acid identities between *Desulfosporosinus* and *Desulfitobacterium* species genomes (in%, written into cells). The dendrogram is based on Fig. S2b.

**Fig. S4.**
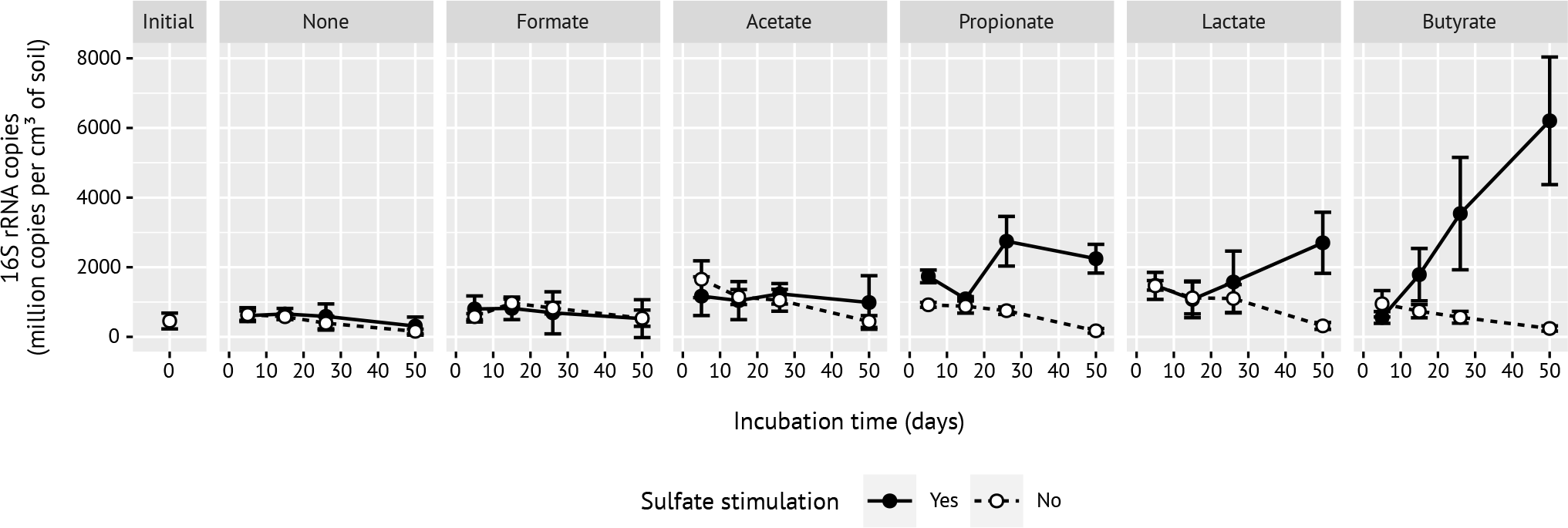
Time-resolved 16S rRNA copies of the low-abundance *Desulfosporosinus* population as determined by quantitative PCR, modified from (7). Error bars are ± one standard deviation (n=3; n=2 for propionate with sulfate stimulation, all days, and butyrate with sulfate stimulation, day 50).(7). Solid lines and symbols represent sulfate-stimulated microcosms whereas dashed lines and open symbols represent control microcosms without external sulfate. Panels represent the various substrate incubations, initial stands for initial peat soil.

**Fig. S5.**
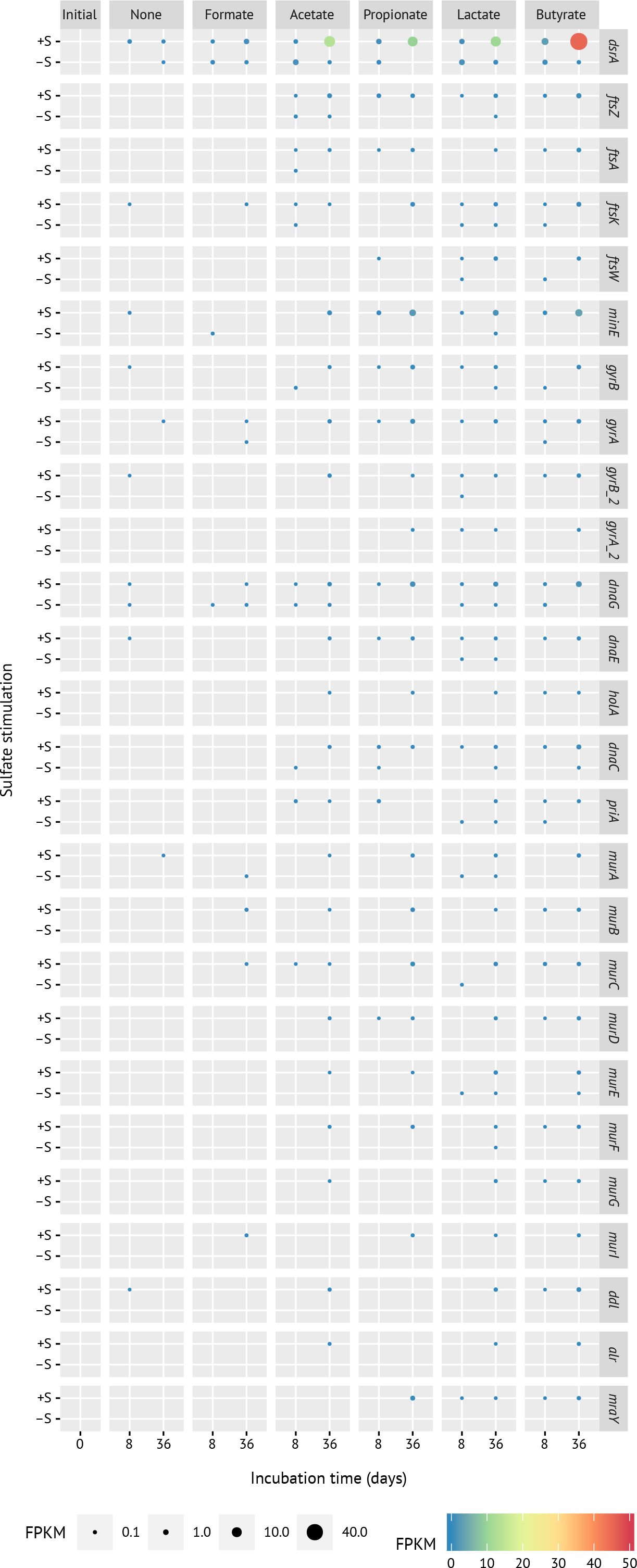
Time-resolved changes of all unambiguously identified genes related to cell division (*ftsZ, ftsA, ftsK, ftsW, minE*), DNA replication (*gyrB, gyrA, dnaG, dnaE, holA, dnaC, priA*), and cell envelope biogenesis (*murABCDEFGI, ddl, alr, mraY*, Table S1); *dsrA* is included as reference, analogous to Fig. 3. Panels represent the various substrate incubations: initial, initial peat soil to set up peat microcosms; +/−S, incubations with or without external sulfate. The size and color of the dots represent average FPKM values of the respective normalized gene expression.

